# The phosphatase inhibitor LB-100 acts synergistically with the NPR2 agonist BMN-111 to improve bone growth

**DOI:** 10.1101/2020.09.10.288589

**Authors:** Leia C. Shuhaibar, Nabil Kaci, Jeremy R. Egbert, Léa Loisay, Giulia Vigone, Tracy F. Uliasz, Emilie Dambroise, Mark R. Swingle, Richard E. Honkanen, Laurinda A. Jaffe, Laurence Legeai-Mallet

**Affiliations:** Department of Cell Biology, University of Connecticut Health Center, Farmington CT 06030 USA; Université de Paris, Imagine Institute, Laboratory of Molecular and Physiopathological Bases of Osteochondrodysplasia. INSERM UMR 1163, F-75015, Paris, France; Department of Biochemistry and Molecular Biology, University of South Alabama, Mobile AL 36688, USA

## Abstract

Activating mutations in fibroblast growth factor receptor 3 (FGFR3) and inactivating mutations in the natriuretic peptide receptor 2 (NPR2) guanylyl cyclase both result in decreased production of cyclic GMP (cGMP) in chondrocytes and severe short stature, causing achondroplasia (ACH) and acrosomelic dysplasia type Maroteaux, respectively. Previously we showed that an NPR2 agonist BMN-111 (vosoritide) increases bone growth in mice mimicking ACH (*Fgfr3*^*Y367C/+*^), and that in control growth plate chondrocytes, FGFR3 signaling decreases NPR2 activity by dephosphorylating the NPR2 protein. Here we tested whether a phosphatase inhibitor (LB-100) could enhance bone growth in ACH. In *ex vivo* imaging experiments using a FRET sensor to measure cGMP production in chondrocytes of living tibias from newborn mice, LB-100 counteracts the FGF-induced dephosphorylation and inactivation of NPR2. In *ex vivo* experiments with *Fgfr3*^*Y367C/+*^ mice, LB-100 in combination with BMN-111 increases the rate of femur growth by ∼25% vs BMN-111 alone, restores chondrocyte terminal differentiation, increases the proliferative growth plate area of the femur, and reduces the activity of the MAP kinase pathway. Our results provide a proof of concept that a phosphatase inhibitor could be used together with an NPR2 agonist to enhance cGMP production as a therapy for ACH.

**GRAPHICAL ABSTRACT:** 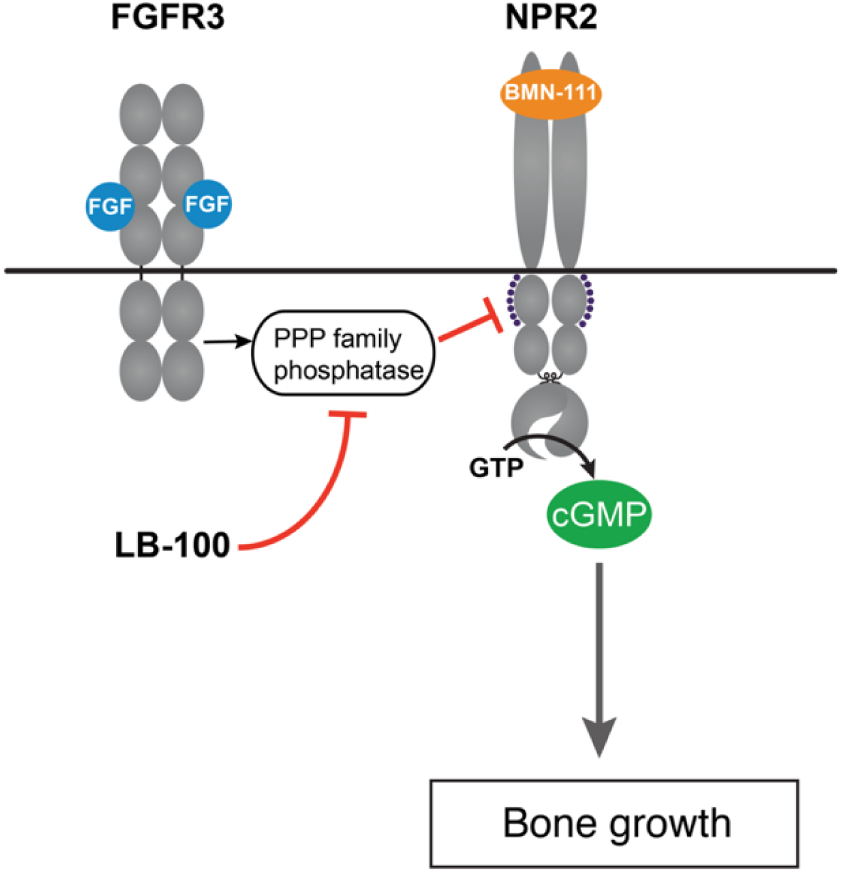

## INTRODUCTION

Achondroplasia (ACH), the most common form of dwarfism, is due to a gain of function mutation in the fibroblast growth factor receptor type 3 (*FGFR3*) gene (1,2). FGFR3 is expressed in growth plate cartilage and bone, which explains the bone anomalies observed in patients with ACH. The characteristic features of these patients are short arms and legs, macrocephaly, hypoplasia of the midface, lordosis, spinal stenosis, and low bone mineral density (3). The generation of *Fgfr3*-specific mouse models has highlighted the role of FGFR3 during bone growth. In the absence of *Fgfr3*, the most prominent phenotype of the mice is overgrowth, thus indicating that FGFR3 is a negative regulator of bone growth (4,5). Conversely, mice expressing a *Fgfr3*-activating mutation develop dwarfism, and have reduced linear growth and impaired endochondral ossification with reduced chondrocyte proliferation and reduced hypertrophic differentiation (6-9). A complex intracellular network of signals including FGFR3 mediates this skeletal phenotype. Activating mutations in FGFR3 lead to upregulated FGFR3 protein (10) and to increased activity of several downstream intracellular signaling pathways including MAPK, PI3K/AKT, PLCγ and STATs (11).

During development, the rate of longitudinal bone growth is determined by chondrocyte proliferation and differentiation and is regulated by several secreted growth factors and endocrine factors, including parathyroid hormone-like peptide, Indian Hedgehog, bone morphometric proteins, transforming growth factor β, insulin like growth factor, and C-type-natriuretic peptide (CNP) (12). CNP and its receptor, the guanylyl cyclase natriuretic peptide receptor 2 (NPR2), are expressed in chondrocytes as well as in osteoblasts and are recognized as important regulators of longitudinal bone growth and bone homeostasis. NPR2 possesses guanylyl cyclase activity that leads to synthesis of cyclic guanosine-3’,5’-monophosphate (cGMP), and dysregulation of this pathway is responsible for skeletal disorders. In clinical studies, inactivating mutations of NPR2 were found to cause a rare form of extreme short stature, called acromesomelic dysplasia type Maroteaux (13-18). Conversely, heterozygous NPR2 gain of function mutations cause tall stature (19) and overexpression of CNP due to a balanced translocation is responsible for overgrowth and bone anomalies (20,21). Animal models with *Npr2* loss of function mutations or with disruption of the CNP gene (*Nppc*) show severe dwarfism, while overstimulation of CNP and NPR2 causes overgrowth disorders. All of these data support a key role of the CNP/NPR2 signaling pathway for normal growth (22,23).

Our recent studies have indicated that among its diverse signaling effects, activation of FGFR3 results in reduced phosphorylation and activity of NPR2 in the growth plate, thus lowering cGMP and opposing bone growth (24,25). Mechanistically, the CNP-induced increase in bone growth is due at least in part to cGMP counteracting the FGF-induced decrease in chondrocyte cell division by inhibiting the ERK pathway that is stimulated by FGF (26,27). Synthesis of cGMP by NPR2 requires extracellular binding of CNP (28), and CNP or a hydrolysis-resistant CNP analog, known as BMN-111 or vosoritide, increases bone growth in mouse models of achondroplasia, demonstrating a significant recovery of bone growth mediated by NPR2/cGMP signaling (29,30). These preclinical studies were confirmed in humans, and vosoritide is currently in clinical development, with phase two results showing additional height gain in achondroplasia patients treated with vosoritide (31). CNP activation of NPR2 activity requires that the receptor is also phosphorylated on multiple serines and threonines (28,32). A PPP-family phosphatase mediates the FGF-induced dephosphorylation and inactivation of NPR2, suggesting that a phosphatase inhibitor could enhance bone growth if applied together with CNP (24,25).

Here we tested this concept using a semi-selective PPP family phosphatase inhibitor, LB-100 (33). In studies of animal cancers, LB-100 has been shown to enhance responses to immunotherapy, CAR-T cell therapy, and tyrosine kinase inhibitors (34-36). Phase one clinical trials concluded that the safety, tolerability and preliminary evidence of anti-tumor activity supported continual testing as a novel treatment for human cancers (37). Here we find that LB-100 counteracts the FGF-induced dephosphorylation and inactivation of NPR2, complementing the CNP stimulation and promoting bone growth in a mouse model of achondroplasia. Our results provide evidence for the concept that an inhibitor of NPR2 dephosphorylation could be could used together with an NPR2 agonist to enhance cGMP production as therapy for ACH.

## RESULTS

### LB-100 counteracts the inactivation of NPR2 by FGF in growth plate chondrocytes

NPR2 activity in chondrocytes of intact growth plates was measured as previously described, using mice expressing a FRET sensor for cGMP, cGi500, and wildtype *Fgfr3* (25). Tibias were isolated from newborn mice, and the overlying tissue was excised to expose the growth plate for confocal imaging (***Figure 1A***). When the NPR2 agonist CNP was perfused across the growth plate, the CFP/YFP emission ratio from cGi500 increased, indicating an increase in cGMP, due to stimulation of the guanylyl cyclase activity of NPR2 (***Figure 1B***). Perfusion of A-type natriuretic peptide (ANP), which activates the NPR1 guanylyl cyclase, or perfusion of a nitric oxide donor (DEA/NO), which activates soluble guanylyl cyclases, did not increase cGMP (***Figure S1***), showing that among the several mammalian guanylyl cyclases, only NPR2 is active in the chondrocytes of the mouse growth plate. As previously shown (25), exposure of the growth plate to FGF18 suppressed the cGMP increase in response to CNP perfusion (***Figure 1B***), indicating that FGF receptor activation decreases NPR2 activity.

**Figure 1.**
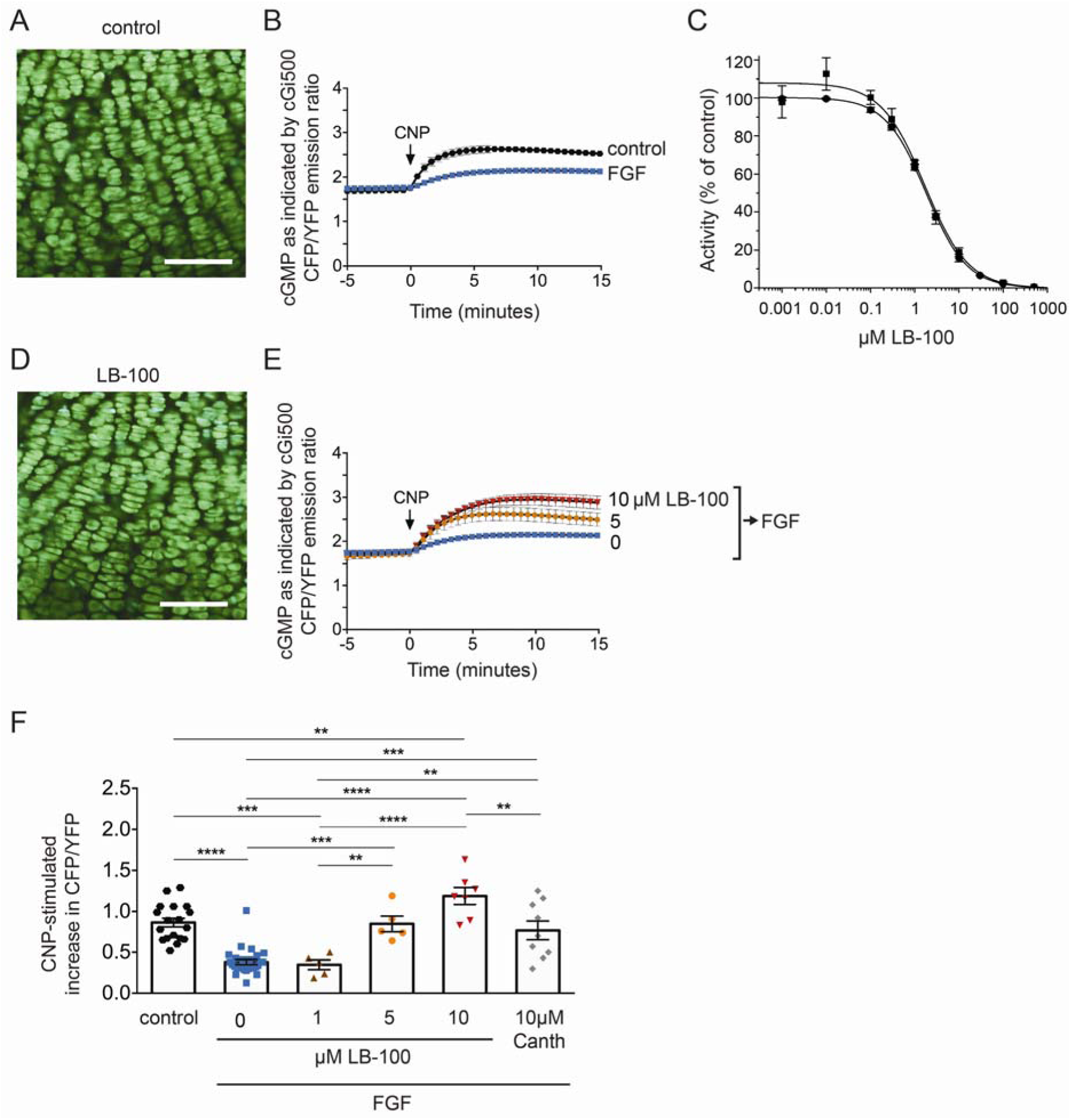
LB-100 counteracts the inactivation of NPR2 by FGF in growth plate chondrocytes of intact tibias from newborn mice. (**A**) Confocal image of cGi500 fluorescence in chondrocytes in the columnar and prehypertrophic region of the tibia growth plate, as used for measurements of cGMP production. Scale bar = 100 µm. (**B**) cGMP increase in growth plate chondrocytes in response to perfusion of 0.1 µM CNP, and inhibition of the cGMP increase by pretreatment of the tibias with 0.5 µg/ml FGF18 + 1 µg/ml heparin for 80 minutes before starting the recording. The control tibias were pretreated for 80 minutes with heparin only. The graph shows the mean ± SEM for 27 control tibias and 18 FGF-treated tibias. (**C**) Inhibitory effect of LB-100 on the activity of PPP1CA by LB-100 assayed using DiFMUP (diamonds) or [^32^P]-labeled histone (squares) as a substrate. Assays were conducted as described in the Methods. Each point represents the mean +/- SD (n=3-4). IC_50_ values are provided in Table 1. (**D**) Confocal image of cGi500 fluorescence in growth plate chondrocytes after pretreatment of the tibia with 10 µM LB-100 for 2 hours. No difference in morphology was seen compared with control tibias (A) without LB-100. Scale bar = 100 µm. (**E**,**F**) Effect of LB-100 (or cantharidin) preincubation on CNP-stimulated cGMP production in FGF-treated tibias. Tibias expressing cGi500 were preincubated with solutions with or without LB-100 for 60 minutes. FGF was then added, and 80 minutes later, tibias were placed in a perfusion slide for cGi500 imaging during CNP perfusion. (**E**) shows the CFP/YFP emission ratio as a function of time after CNP perfusion. (**F**) shows the CFP/YFP emission ratio at 15 minutes after CNP perfusion. Symbols indicate individual tibias (n = 5-27). For **E** and **F**, bars show mean ± SEM. Data were analyzed by one-way ANOVA followed by the Holm-Sidak multiple comparison test (**p < 0.01, ***p < 0.001, **** p < 0.0001).

Based on previous evidence that a PPP-family phosphatase inhibitor, cantharidin (100 µM), inhibits the inactivation of NPR2 in growth plate chondrocytes by FGF (25), we tested whether a less toxic cantharidin derivative, LB-100, would increase NPR2 activity and long bone growth. LB-100 was originally reported as a specific inhibitor of PP2A (PPP2CA) but was later shown to also act as a catalytic inhibitor of PPP5C (33). As cantharidin demonstrates only modest selectivity for PPP2CA versus PPP1C (38), we tested the ability of LB-100 to inhibit PPP1C activity using two established assays that use different substrates. We observed that LB-100 also inhibits PPP1C with an IC_50_ < 2 μM (***Figure 1C, Table 1***). Based on its structural similarity with cantharidin, 10 µM LB-100 is not likely to inhibit PPP3C/calcineurin, PPP4C, or PPP7C/PPPEF (39).

**Table 1.** Inhibitory activity of LB-100 against PPP1CA, PPP2CA, and PPP5C

| Substrate | Phosphatase<br>IC<sub>50</sub> mean ± SEM (μM) | | |
| --- | --- | --- | --- |
|  | PPP1CA | PPP2CA | PPP5C |
| DiFMUP | 1.80 ± 0.02 | 0.39 ± 0.013 | 1.82 ± 0.093 |
| [<sup>32</sup>P]-labeled phosphohistone | 1.68 ± 0.13 | 0.64 ± 0.037 | 4.9 ± 0.29 |

To investigate if LB-100 counteracts the inactivation of NPR2 by FGF, we preincubated the tibia with or without LB-100 and then with FGF. Following these incubations, the tibia was placed in a perfusion slide for confocal imaging, and cGMP production by NPR2 was monitored by measuring the increase in CFP/YFP emission ratio in response to CNP. Incubation with 10 µM LB-100 caused no visible change in chondrocyte morphology as imaged in the live growth plate, indicating no obvious toxicity (compare ***Figure 1D*** to the control in ***Figure 1A***).

After FGF treatment, the cGMP increase in response to CNP was small (***Figure 1E***). However, when the tibia was preincubated with 5 or 10 µM LB-100 before applying FGF, the CNP-induced cGMP increase was enhanced (***Figure 1E,F***). 1 µM LB-100 had no effect (***Figure 1F***). The CFP/YFP emission ratio attained after CNP perfusion in tibias that had been incubated in 5 or 10 µM LB-100 before the FGF treatment was similar to or greater than that in control tibias without FGF (***Figure 1F***). ***Figure 1F*** summarizes the CNP-stimulated increases in the CFP/YFP emission ratio from cGi500 under these various conditions, and demonstrates that LB-100 counteracts the inactivation of NPR2 by FGF. LB-100 was more effective than cantharidin, with 5 µM LB-100 resulting in a stimulation equivalent to that seen with 10 µM cantharidin (***Figure 1F***).

### LB-100 counteracts the FGF-induced dephosphorylation of NPR2 by FGF in primary chondrocyte cultures

To investigate if LB-100 counteracts the FGF-induced dephosphorylation of NPR2, we used Phos-tag gel electrophoresis (40) to analyze the phosphorylation state of NPR2 in isolated chondrocytes from the ribs of newborn mice. After 4 days in culture, the chondrocytes had formed a monolayer with a mosaic appearance (***Figure 2A***). A 1 hour incubation with 10 µM LB-100 did not cause any visible change in cell morphology (***Figure 2B***).

**Figure 2.**
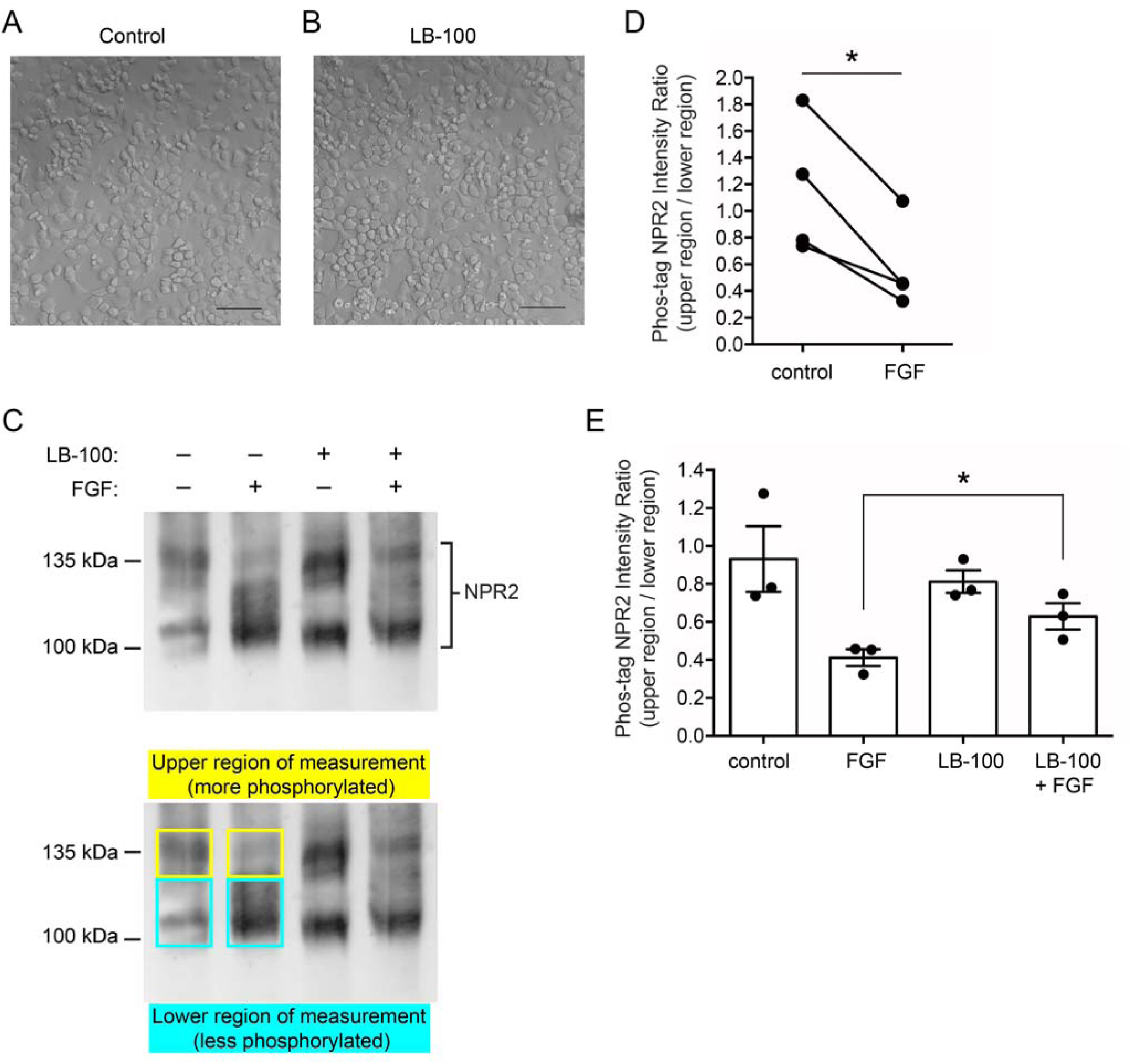
LB-100 counteracts the FGF-induced dephosphorylation of NPR2 in primary chondrocyte cultures. (**A, B**) Images of chondrocytes isolated from newborn mouse ribs, cultured for 4 days and treated without or with LB-100 (10 µM) for 1 hour. Scale bars = 100 µm. (**C**) Western blot of a Phos-tag gel showing the decrease in NPR2 phosphorylation in chondrocytes treated with FGF18 (0.5 µg/ml) for 10 minutes, and attenuation of this dephosphorylation in chondrocytes pretreated with LB-100 (10 µM) for 1-hour prior to treatment with FGF18. The lower panel shows the regions used for densitometry measurements. (**D**) Densitometry measurements comparing chondrocytes treated with or without FGF18. The y-axis indicates the ratio of the intensity of the upper region to that of the lower region as shown in C; a smaller ratio indicates a decrease in NPR2 phosphorylation (measurements for 4 experiments similar to that shown in C; analyzed by paired t-test). (**E**) Densitometry measurements comparing chondrocytes treated with or without LB-100, then with or without FGF18 (mean ± SEM for the 3 experiments shown in D for which LB-100 was also tested; the two indicated groups were analyzed by paired t-test). Asterisks indicate significant differences (p<0.05).

We then compared the phosphorylation state of NPR2 in chondrocytes with and without LB-100 preincubation, and with and without subsequent exposure to FGF. Chondrocyte proteins were separated by Phos-tag gel electrophoresis, which retards migration of phosphorylated proteins, and western blots were probed for NPR2 (***Figure 2C***). Without FGF treatment, NPR2 protein from the rib chondrocytes was present in a broad region of the gel. With FGF treatment, the ratio of the signal in the upper vs lower regions decreased (***Figure 2C,D***), indicating NPR2 dephosphorylation in response to FGF and confirming, with primary chondrocytes, a previous study using a rat chondrosarcoma (RCS) cell line (24). However, if the chondrocytes were preincubated with 10 µM LB-100, the dephosphorylation in response to FGF was only partial, indicating that LB-100 counteracts the FGF-induced dephosphorylation of NPR2 (***Figure 2C,E***). Similar results were obtained by probing blots of Phos-tag gels with a different antibody recognizing NPR2 (***Figure S2***).

### In *Fgfr3*^*Y367C/+*^ femurs, LB-100 enhances the stimulation of bone growth by the hydrolysis resistant NPR2 agonist BMN-111

Previously we showed that the hydrolysis-resistant CNP analog BMN-111 increases bone growth in a mouse model of achondroplasia in which tyrosine 367 is changed to a cysteine (*Fgfr3*^*Y367C/+*^), resulting in constitutive activation of FGFR3 (30,41). However, BMN-111 only partially rescued the effect of the FGFR3-activating mutation. Our finding that LB-100 opposes the FGF inhibition of NPR2 activity in chondrocytes suggested that applying LB-100 together with BMN-111 might enhance the stimulation of growth (***Figure 3A***). Like CNP, BMN-111 (0.1 µM) stimulated NPR2 activity in growth plate chondrocytes (***Figure S3***).

**Figure 3.**
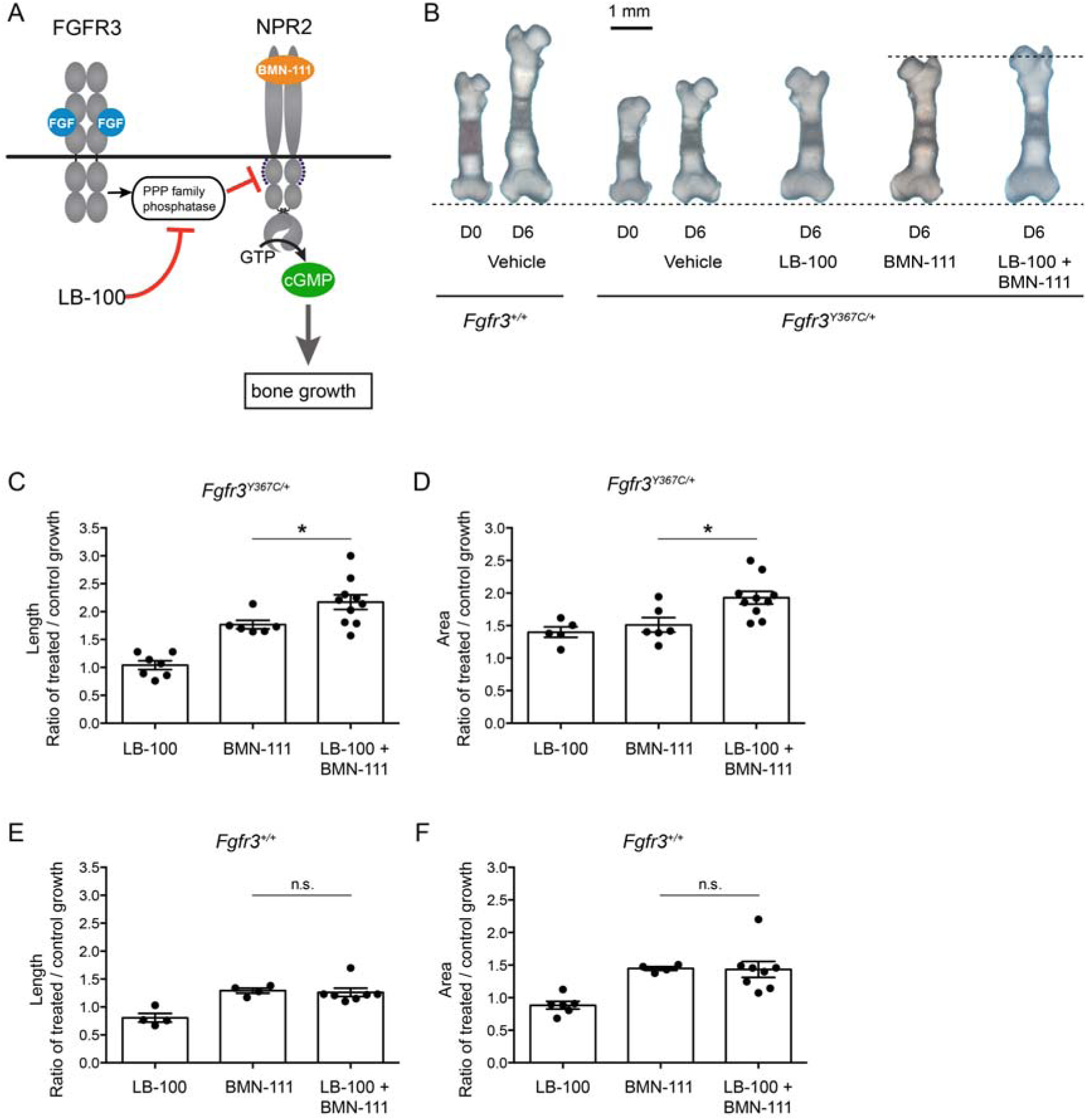
LB-100 and BMN-111 act synergistically to stimulate growth in fetal femurs from *Fgfr3*^*Y367C/+*^ mice. (**A**) Diagram showing sites of action of LB-100 and BMN-111. (**B**) Representative photographs of fetal femurs from E16.5 day old *Fgfr3*^+/+^ (wildtype) and *Fgfr3*^*Y367C/+*^ mice, before (D0) and after a 6 day (D6) culture with the indicated treatments. The upper dashed line indicates the groups compared in C-F. (**C**,**D**) Measurements of growth in bone length (**C**) and area (**D**), showing that in *Fgfr3*^*Y367C/+*^ bones, BMN-111+LB-100 increases growth more than BMN-111 alone. (**E**,**F**) Measurements of growth in bone length (**E**) and area (**F**) for *Fgfr3*^+/+^ bones, showing that for *Fgfr3*^+/+^ bones, BMN-111+LB-100 does not increase growth more than BMN-111 alone. For all experiments, the concentration of BMN-111 was 0.1 µM, and the concentration of LB-100 was 10 µM. Symbols in graphs C-E indicate individual bones (n = 4-10). Bars represent mean ± SEM and data were analyzed by unpaired two-tailed t-tests. Asterisks indicate significant differences (p < 0.05) between indicated groups; n.s. indicates no significant difference (p > 0.05).

As previously reported (30), 0.1 µM BMN-111 increased the rate of elongation of cultured femurs from embryonic day 16.5 *Fgfr3*^*Y367C/+*^ mice (***Figures 3 B,C, S4***). The mean rate of increase in bone length in the BMN-111-stimulated *Fgfr3*^*Y367C/+*^ femurs was 1.77 times that in vehicle-treated bones (***Figure 3C***). LB-100 alone did not significantly increase the rate of bone elongation, showing a growth rate ratio of 1.04 for LB-100/control (***Figure 3C***). The fact that LB-100 alone, without BMN-111, does not affect bone elongation is consistent with phosphorylation of NPR2 not affecting basal (non-CNP-dependent) activity of NPR2 (42). However, when *Fgfr3*^*Y367C/+*^ femurs were cultured with BMN-111 together with 10 µM LB-100, the mean rate of bone elongation increased to 2.17 times that in untreated bones (***Figure 3C***). Thus, the combination of BMN-111 and LB-100 significantly increased the bone elongation rate to a level ∼23% higher than the level observed with BMN-111 alone.

In addition to these measurements of bone length, we also measured the effect of LB-100 and BMN-111 on the rate of increase of the total bone and cartilage area, defined as the area within the periphery of a photograph of the femur (see ***Figure S4***). Based on these area measurements, LB-100 by itself stimulated growth of femurs from *Fgfr3*^*Y367C/+*^ mice, as did BMN-111 by itself, with a growth rate ratio of 1.40 for LB-100/control, and a growth rate ratio of 1.51 for BMN-111/control (***Figure 3D***). The combination of LB-100 and BMN-111 was even more effective, with a growth rate ratio of 1.93. Thus, the combination of BMN-111 and LB-100 enhanced the rate of increase in area by 27% compared with BMN-111 alone (***Figure 3D***).

With cultured femurs from *Fgfr3*^*+/+*^ mice, BMN-111 increased the rate of bone growth, but combining BMN-111 with LB-100 did not enhance the growth rate beyond that seen with BMN-111 alone (***Figure 3E,F***). This result contrasts with the ability of LB-100 to enhance BMN-111-stimulated bone growth in *Fgfr3*^*Y367C/+*^ mice. Because *Fgfr3*^*Y367C/+*^ mice have elevated FGFR3 tyrosine kinase activity due to an activating *Fgfr3* mutation (43), their NPR2 would be expected to be less phosphorylated and less active, and LB-100 would be expected to restore their NPR2 phosphorylation and activity towards *Fgfr3*^*+/+*^ levels (see ***Figures 1 and 2***), thus increasing bone growth. In contrast, if baseline NPR2 phosphorylation is higher in *Fgfr3*^*+/+*^ vs *Fgfr3*^*Y367C/+*^ mice, the growth stimulating effect of LB-100 might be less.

### Combined treatment with LB-100 and BMN-111 improves growth plate cartilage homeostasis

Histological analyses of the epiphyseal growth plates of *Fgfr3*^Y367C/+^ femurs showed that combining BMN-111 and LB-100 treatment modified cartilage growth homeostasis (***Figure 4***). Prehypertrophic and hypertrophic chondrocytes produce an extracellular matrix rich in Collagen type X (COLX), allowing us to use COLX immunostaining to label the hypertrophic region, and to visualize and measure individual cells. This labeling revealed a highly beneficial effect of the combined treatment on the size of the cells in the hypertrophic area of *Fgfr3*^Y367C/+^ femurs (***Figure 4***). The mean cross-sectional area of the hypertrophic chondrocytes in the proximal growth plate of *Fgfr3*^Y367C/+^ mice was reduced by about half compared to that in the *Fgfr3*^+/+^ growth plate (***Figures 4A,B; S5)***. As previously reported (30), BMN-111 increased the size of the *Fgfr3*^Y367C/+^ hypertrophic chondrocytes, but the cells were smaller than for the wildtype (***Figure 4A,B***). However, with the combined treatment of BMN-111 and LB-100, the mean area of the *Fgfr3*^*Y367C/+*^ hypertrophic cells in the proximal growth plate was 32% greater than with BMN-111 alone, and similar to that of *Fgfr3*^*+/+*^ hypertrophic cells, indicating that the final differentiation of the chondrocytes was restored by the treatment (***Figure 4A,B***). Corresponding measurements for the distal growth plate showed a similar trend, but the difference for bones treated with BMN-111 vs BMN-111 + LB-100 was not statistically different (***Figure S6***).

**Figure 4.**
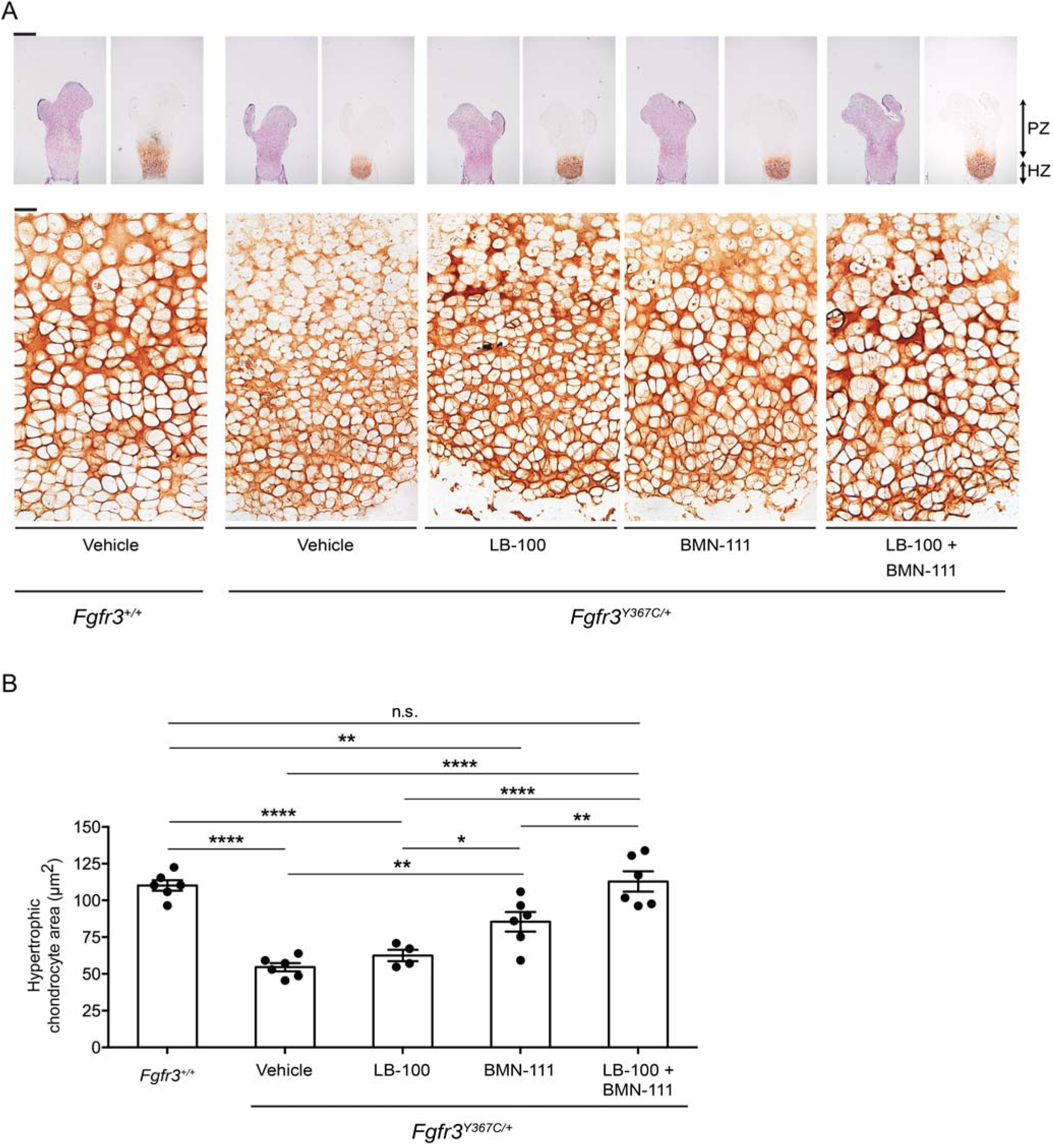
Dual action of LB-100 and BMN-111 improves chondrocyte differentiation in growth plates of *ex-vivo* cultured *Fgfr3*^*Y367C/+*^ femurs. (**A**) Representative images of HES-stained and type X collagen immunostained proximal growth plates of embryonic *Fgfr3*^*Y367C/+*^ femurs incubated for 6 days with vehicle or LB-100 (10 µM) and/or BMN-111 (0.1 µM). Growth plates of *Fgfr3*^*+/+*^ femurs cultured with vehicle are also shown. PZ = proliferative zone. HZ = hypertrophic zone. Scale bars: 400 μm for upper row, 20 µm for lower row. (**B**) Mean area of individual hypertrophic chondrocytes in proximal growth plates of femurs treated as described for A (n = 4-6 bones measured for each condition, with 54-137 cells measured for each bone). Data were analyzed by one-way ANOVA followed by the Holm-Sidak multiple comparison test (*p < 0.05, **p < 0.01, ****p<0.0001).

We also observed a beneficial effect of the combined treatment on the proliferative region of the growth plate. We measured the area of the proliferative region by subtracting the hypertrophic area, identified by COLX labeling, from the total growth plate area. Based on these measurements, the combined treatment increased the total proliferative growth plate area of the femur by an average of 33% over vehicle, compared to 20% for BMN-111 alone (***Figure S7)***. Thus, the combined treatment increased the proliferative area by 13% compared to BMN-111 alone (***Figure S7)***. In summary, the combined treatment both increased the proliferative growth plate area of the femur and restored chondrocyte terminal differentiation.

CNP signaling through NPR2 in the growth plate inhibits the MAP kinase pathway and its extracellular signal-regulated kinase 1 and 2 (ERK1/2) (26,27). Therefore, we investigated the impact of treatment with LB-100 + BMN-111 on the phosphorylation of ERK1/2. In agreement with the constitutive activity of the Y367C *Fgfr3* gain-of-function mutation acting to decrease NPR2 activity (***Figure 5A***), immunolabeling showed a high level of phosphorylated ERK1/2 in the proximal and distal parts of the cartilage compared to wildtype controls (***Figure 5B,C***). The combined LB-100 and BMN-111 treatment of *Fgfr3*Y^367C/+^ femurs decreased the activity of the MAP kinase pathway as demonstrated by the decreased phosphorylation of ERK1/2 in the proximal and distal growth plates of the femurs (***Figure 5B,C***). These data are in agreement with 1) the role of the MAP kinase pathway as a regulator of chondrocyte differentiation and with 2) our hypothesis that the elevation of cGMP inhibits the MAP kinase pathway thus promoting bone growth.

**Figure 5.**
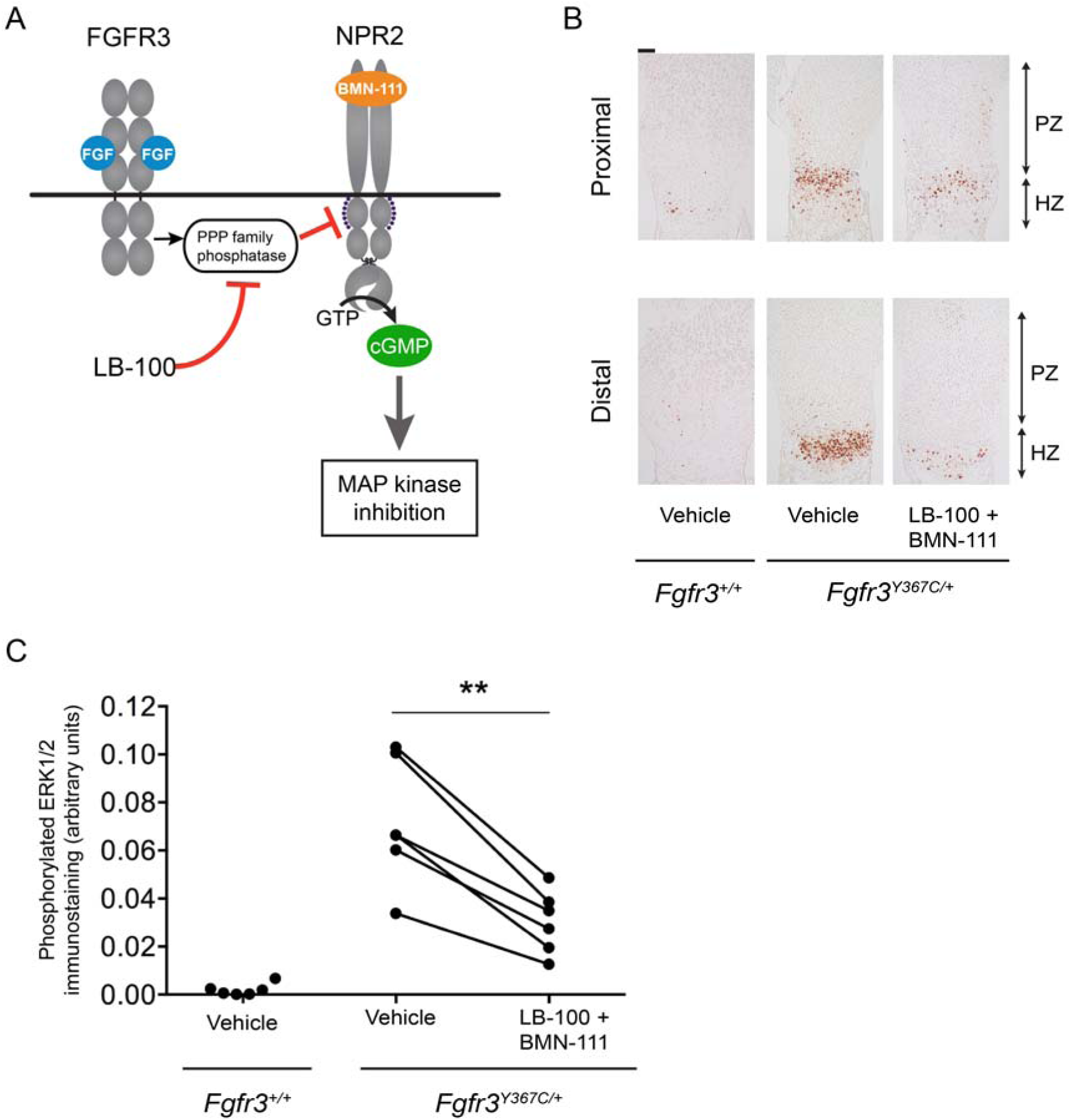
Dual action of LB-100 and BMN-111 decreases the activation of the MAP kinase pathway. (**A**) Diagram showing sites of action of LB-100 and BMN-111 on the MAP kinase pathway. (**B**) Representative images of phosphorylated ERK1/2 immunostaining of proximal and distal growth plates of embryonic Fgfr3^Y367C/+^ femurs incubated for 6 days with vehicle or with LB-100 (10 µM) and BMN-111 (0.1 µM). Growth plates of wildtype (Fgfr3^+/+^) femurs cultured with vehicle are also shown. Scale bar: 100 µm. (**C**) Quantitation of phosphorylated ERK1/2 immunostaining in proximal plus distal growth plates of femurs from the same animal treated with either vehicle or a combination of LB-100 and BMN-111. Data were analyzed by paired t-test; ** indicates p < 0.01. Data from vehicle-treated *Fgfr3*^+/+^ femurs are shown for comparison.

## DISCUSSION

Understanding of the mechanisms used by FGFR3 and CNP as important regulators of longitudinal bone growth has allowed the development of an effective therapeutic strategy using a CNP analog (vosoritide, also known as BMN-111) to treat achondroplasia (31). The findings described here identify the PPP-family phosphatase inhibitor LB-100 as a stimulator of bone growth when used in combination with this CNP analog to stimulate production of cGMP by NPR2. Firstly, using isolated bones incubated with FGF to mimic an achondroplasia-like condition, we show that pretreatment with LB-100 counteracts the decrease in NPR2 guanylyl cyclase activity by FGFR3. Secondly, our results support the hypothesis that FGFR3 activation leads to NPR2 dephosphorylation in chondrocytes and that LB-100 suppresses the dephosphorylation. Moreover, application of a combination of BMN-111 and LB-100 to bones from the achondroplasia mouse model *Fgfr3*^*Y367C/+*^ results in growth that exceeds that stimulated by BMN-111 alone, providing a proof of concept that BMN-111 and a PPP-family phosphatase inhibitor could potentially be used in combination for treatment of skeletal dysplasias such as achondroplasia.

Our data also show the benefit of this treatment for growth plate cartilage during bone development in *Fgfr3*^*Y367C/+*^ mice. During the process of endochondral ossification, chondrocytes actively proliferate in the resting and proliferating chondrocyte zone, and then differentiate to hypertrophic chondrocytes, which lose the capacity to proliferate. The terminally differentiated hypertrophic cells are removed by cell death or transdifferentiate into osteoblasts. It is well known that FGFR3 signaling decreases bone growth by inhibiting both chondrocyte proliferation and differentiation and bone formation, and it has been proposed that FGFR3 acts by way of ERK1/2 to restrict hypertrophic differentiation (44). Here, we showed that the treatment with BMN-111 and LB-100 reduced the levels of phosphorylated ERK1/2, thus modifying chondrocyte differentiation and allowing bone growth. In addition, we noted an impressive increase in the size of the hypertrophic cells. We concluded that the treatment perfectly restored cartilage homeostasis, and we hypothesize that the elevated cGMP resulting from this treatment could be a key regulator of transdifferentiation of hypertrophic cells into osteoblasts and could control the chondrogenic or osteogenic fate decision.

The increase in NPR2 phosphorylation by LB-100 is correlated with improved chondrocyte proliferation in *Fgfr3*^*Y367C/+*^ femurs, consistent with previous results with a mouse model (*Npr2*-7E) mimicking constitutive phosphorylation of NPR2 (25). Because LB-100 inhibits multiple PPP-family phosphatases (33, and ***Table 1***), and because its safety for long term use in children is unknown, our results provide only a proof of principle for a possible combination treatment. Future studies to determine which phosphatase(s) acts to dephosphorylate NPR2 in chondrocytes are clearly warranted, and the development of more specific inhibitors targeting the responsible phosphatase(s) could lead to future therapies.

Recent mouse studies indicate that in addition to increasing prepubertal bone elongation, phosphorylation of NPR2 increases bone density, due to an increase in the number of active osteoblasts at the bone surface (45). Because low bone density is one of the key clinical features of achondroplasia (46), the combination a CNP analog and a phosphatase inhibitor could also have a beneficial impact on bone density for patients with achondroplasia and related conditions. In addition, such a treatment could have potential for treatment of osteoporosis, and because CNP/NPR2 also plays a key role in regulation of joint homeostasis, could be beneficial for preventing or minimizing cartilage loss and promoting repair of the damaged articular cartilage in skeletal disorders and osteoarthritis (47). More generally, the combination of natriuretic peptides and phosphatase inhibitors could have therapeutic potential for multiple disorders involving NPR2 and the related guanylyl cyclase NPR1 that also requires phosphorylation for activity (48).

In summary, the combined (LB-100 + BMN-111) treatment acts on both chondrocyte proliferation and differentiation, thus promoting better bone growth. In achondroplasia, the homeostasis of the growth plate is disturbed, and proliferation and differentiation are affected by the overactivation of FGFR3. Currently, BMN-111 (vosoritide) is being studied in children with ACH, and as demonstrated in preclinical studies, mostly restores the defective differentiation in the growth plate (30). Recently reported phase 2 data demonstrated that vosoritide results in a sustained increase in annualized growth velocity for up to 42 months in children 5 to 14 years of age with achondroplasia (31). The present study provides a proof of concept that a combination of vosoritide and a phosphatase inhibitor has the potential to increase bone growth rate in ACH patients to a higher level than vosoritide alone.

## METHODS

### Mice

Two mouse lines were used for this study: cGi500 (48) and *Fgfr3*^Y367C/+^ (41). The cGi500 mice were provided by Robert Feil (University of Tübingen). The use of the cGi500 mice for monitoring cGMP levels in the growth plate has been verified by ELISA measurements (25). The genotypes of *Fgfr3*^*Y367C/+*^ and *Fgfr3*^*+/+*^ mice were determined by polymerase chain reaction of tail DNA as previously described (41).

### Reagents

CNP and ANP were obtained from Phoenix Pharmaceutical (012-03 and 005-24, respectively). BMN-111 was synthesized by New England Peptide as a custom order with the following sequence: [Cyc(23,39)]H2N-PGQEHPNARKYKGANKKGLSKGCFGLKLDRIGSMSGLGC-OH, as previously described (30). The purity was >95%. DEA/NO was from Cayman Chemical (82100). LB-100 (3-(4-methylpiperazine-1-carbonyl)-7-oxabicyclo[2.2.1]heptane-2-carboxylic acid) was from Selleck Chemicals (S7537) or MedChem Express (HY-18597). Cantharidin was from Tocris (1548). FGF18 was from PeproTech (100-28), and heparin was from Sigma-Aldrich (H4784). DiFMUP (6,8-Difluoro-4-methyl-7-(phosphonooxy)-2H-1-benzopyran-2-one) was from ThermoFisher (D6567).

### Measurements of cGMP production in tibia growth plates using cGi500

cGMP production in chondrocytes within intact growth plates was measured using tibias dissected from newborn mice (0-1 day old) that globally expressed one or two copies of the cGi500 FRET sensor, as previously described (25). Tibias were dissected and cultured overnight on Millicell organotypic membranes (PICMORG50; Merck Millipore Ltd, Cork, IRL) in BGJb medium (GIBCO, 12591-038) with 0.1% bovine serum albumin (MP Biomedicals, 103700), 100 units/ml of penicillin, and 100 µg/ml of streptomycin. In preparation for imaging, each tibia was slit to remove the tissue overlying the growth plate. Where indicated, the tibia was incubated in LB-100, cantharidin, or control medium, followed by addition of FGF18 (0.5 µg/ml + 1 µg/ml heparin) or control medium containing heparin only. The tibia was then placed in a perfusion slide (ibidi USA, 80186, special order with no adhesive) and the growth plate was imaged on the stage of a confocal microscope, as previously described (25).

### Determination of the effect of LB-100 on PPP1C phosphatase activity

The coding sequence of human PPP1CA was expressed as a maltose binding protein fusion in a BL-21 strain of *Escherichia coli* and purified as previously described (50). Phosphohistone phosphatase assays were performed as previously described (50,51). Briefly, LB-100, at the indicated concentrations, or vehicle control (H_2_O), was added to enzyme/buffer aliquots ∼10 minutes prior to starting assays by the addition of [^32^P]-phosphohistone substrate (to a final assay concentration of 300 nM incorporated phosphate). [^32^P]-phosphohistone was prepared by the phosphorylation of bovine brain histone (Sigma: type-2AS) with cAMP-dependent protein kinase (PKA) in the presence of cAMP and [^32^P]-ATP using established methods (51,52). Phosphatase activity was measured by the quantitation of [^32^P]-labeled orthophosphate liberated from the substrate using established protocols (52). DiFMUP (6,8-Difluoro-4-methylumbelliferyl phosphate)-based inhibition assays were conducted as described (51,52), in a 96 well format using DiFMUP (Invitrogen) (100 µM final assay concentration). IC_50_ values were calculated from a 10-point concentration/dose response curve by a 4-parameter logistic fit of the data, using 3-8 replicates per concentration.

### Rib chondrocyte cultures

Rib cages were dissected from newborn (0-2 day old) mice, and trimmed to remove the skin, spinal cord, and soft tissue around the sternum and ribs. Non-chondrocyte tissue was digested away by incubating the rib cages in 2 mg/ml pronase (Roche, 10165921001) in PBS for one hour in a shaking water bath at 37^°^C, and then incubated in 3 mg/ml collagenase D (Roche, 11088866001) in medium for one hour. After washing, the rib cages were transferred to a dish with fresh collagenase D and incubated for 5-6 hours, with trituration at 2 hours, to release the chondrocytes. The isolated cells were passed through a 40 µm nylon cell strainer (Corning, 431750), resuspended in DMEM/F12 medium (GIBCO,11320-033) with 10% fetal bovine serum (GIBCO 10082-139), 100 units/ml of penicillin, and 100 µg/ml of streptomycin, and plated in 35 mm tissue culture dishes, at a cell density corresponding to one newborn mouse per plate. The cells were cultured for 3 days, at which point they were ∼75-90% confluent, then washed with PBS and incubated in serum-free medium for 18 hours. The cells were then incubated in LB-100 (10 µM), or control medium, followed by addition of FGF18 (0.5 µg/ml + 1 µg/ml heparin) or control medium containing heparin only, as described for ***Figure 2***.

At end of the incubation period, dishes were washed in PBS and cells lysed in 250 µl of 1% SDS containing 10 mM sodium fluoride, 1 µM microcystin-LR (Cayman Chemical, 10007188), and protease inhibitor cocktail (Roche, 04 693 159 001). Protein content was determined by a BCA assay (Pierce, 23225). The protein yield per newborn mouse was ∼200-300 µg.

### Phos-tag gel electrophoresis, and western blotting

Proteins were separated in a Phos-tag-containing gel, as previously described (53), except that chondrocyte lysates (50 µg protein) were used without immunoprecipitation. Phos-tag and protein size markers were obtained from Fujifilm Wako Pure Chemical (AAL-107 and 230-02461, respectively). Blots were probed with an antibody that was made in guinea pig against the extracellular domain of mouse NPR2 (54). This antibody was a gift from Hannes Schmidt (University of Tübingen), and has been previously validated for western blots (24). Note that molecular weight markers are only approximate for Phos-tag gels, and can vary between gels.

### *Ex vivo* culture of fetal femurs

Femurs were cultured *ex vivo*, as described previously (30,55). The left femur was cultured in the presence of LB-100 (10 µM), BMN-111 (0.1 µM), or LB-100 (10 µM) + BMN-111 (0.1 µM), and compared with the vehicle-treated right femur. The bone’s length was measured on day 0 (D0) and day 6 (D6). Images were captured with an Olympus SZX12 stereo microscope and quantified using cellSens software (Olympus). The results were expressed as the increase in femur length or area (D6-D0) in the presence or absence of LB-100, BMN-111, or LB-100+ BMN-111. Bone length and area were measured as shown in ***Figure S4***. To generate the graphs shown in ***Figure 3***, the length or 2-dimensional area on day 0 was subtracted from the length or area on day 6 to calculate the amount of growth. These measurements of growth in drug-treated bones were divided by the mean values from corresponding measurements of control (vehicle-treated) bones; the graphs show the ratio of treated/control growth.

### Histology

Fetal femur (E16.5) explants were fixed in 4% paraformaldehyde, decalcified with EDTA (0.4 M), and embedded in paraffin. Serial 5 μm sections were stained with hematoxylin-eosin-safran (HES) reagent, using standard protocols. For immunohistochemical assessment, sections were labeled with the following antibodies and a Dako Envision Kit: anti-COL X (BIOCYC, N.2031501005; 1:50 dilution), and anti-phosphorylated ERK1-2 (Thr180/Tyr182) (Cell Signaling Technology, #4370; 1:100 dilution). Images were captured with an Olympus PD70-IX2-UCB microscope and quantified using cellSens software.

Mean areas of individual hypertrophic chondrocytes were measured from collagen X labelled sections, within a 166 µm wide x 76 µm high box positioned at 50 µm from mineralization front (***Figure S5***). The measurements were made manually using Fiji software and the freehand selection tool. For analysis of the effect of the drug treatments on the area occupied by proliferative chondrocytes, these cells were identified by their round or columnar shape as seen with HES staining, and by the absence of collagen X labeling. We measured the total area occupied by chondrocytes within the whole growth plate and the area occupied by COLX-positive chondrocytes. The area for proliferating chondrocytes was calculated by subtracting the COLX-positive area from the whole growth plate area.

### Statistics

Data were analyzed using Prism 6 (GraphPad Software). To compare more than two groups, we used one-way ANOVA followed by two-tailed t-tests with the Holm-Sidak correction for multiple comparisons, or two-way ANOVA followed by Sidak’s multiple comparisons tests. Two groups were compared using either paired or unpaired two-tailed t-tests, as indicated in the figure legends.

### Study approval

All experiments were conducted as approved by the animal care committees of the University of Connecticut Health Center and the Imagine Institute/ Paris Descartes University.

## AUTHOR CONTRIBUTIONS

LCS, LAJ and LLM designed the research and wrote the paper. NK performed the ex vivo bone growth experiments. LCS and GV performed the cGMP imaging experiments. JRE and TFU performed the chondrocyte cell culture and Phos-tag analysis. LL and ED performed the immunolabelling quantification. MRS and REH determined LB-100 selectivity for inhibition of PPP family phosphatases. LCS, NK, ED, MRS, and JRE prepared the figures.

## ACKNOWLEDGMENTS

This project received a state subsidy managed by the National Research Agency under the “Investments for the Future” program bearing the reference ANR-10-IAHU-01. It was also supported by the National Institutes of Health (R37HD014939 to LAJ, and R01CA060750 to REH), and by the Fund for Science (to LCS and LAJ). We thank the Imagine Institute’s imaging facility for their help with this work. We also thank Robert Feil for sharing the cGi500 mouse line, Hannes Schmidt for providing the NPR2 antibody, Siu-Pok Yee and Deborah Kaback for mouse colony management, Luisa Lestz, Corie Owen, and Valentina Baena for technical assistance, and Julian Liu, Lincoln Potter and Florence Lorget for helpful discussions.

## SUPPLEMENTARY FIGURES

**Figure S1.**
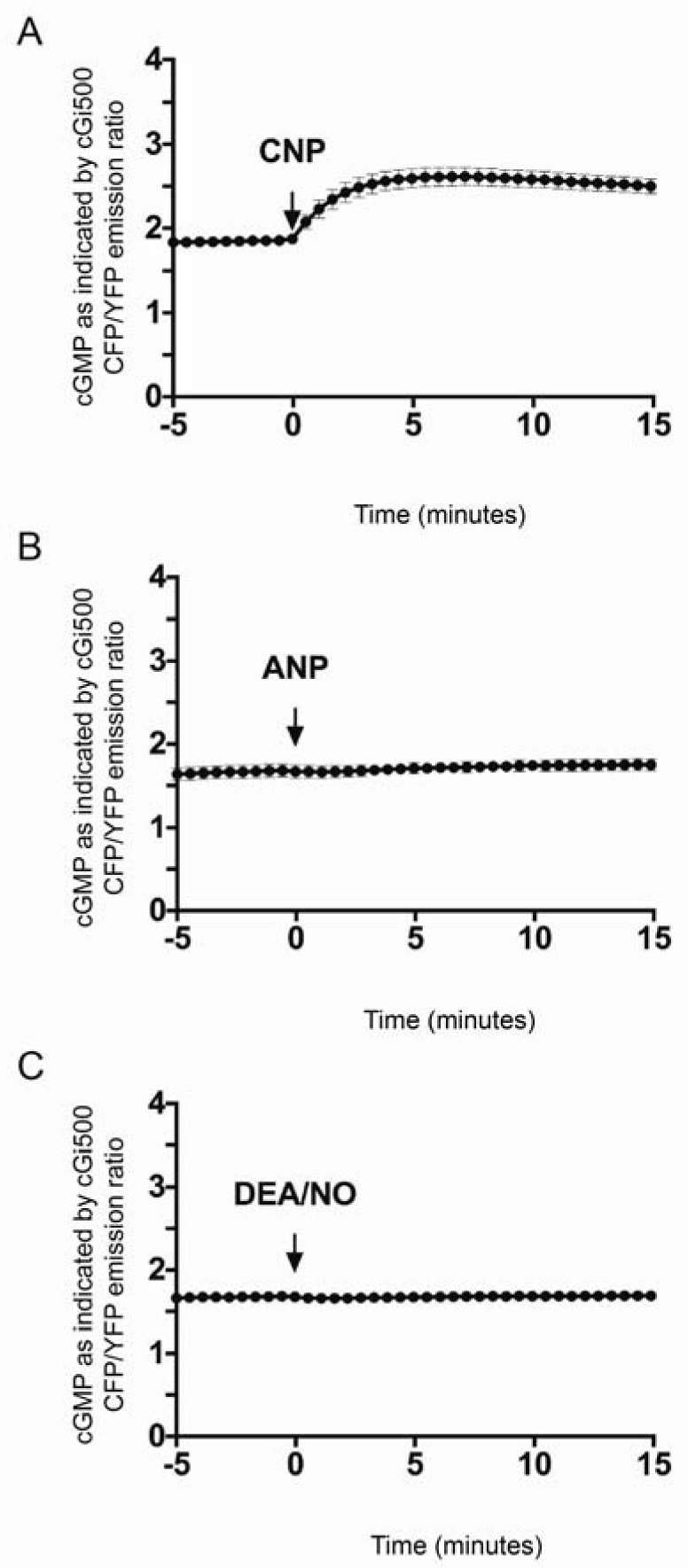
CNP increases cGMP production in growth plate chondrocytes, but ANP and DEA/NO do not. Graphs show the time courses of the CFP/YFP emission ratio from cGi500 after each agonist was perfused across the growth plate. (**A**) 0.1 µM CNP. Mean ± SEM for 27 similar experiments (trace from ***Figure 1B***). (**B**) 0.1 µM ANP. Mean ± SEM for 2 similar experiments. (**C**) 10 µM DEA/NO. Mean ± SEM for 3 similar experiments.

**Figure S2.**
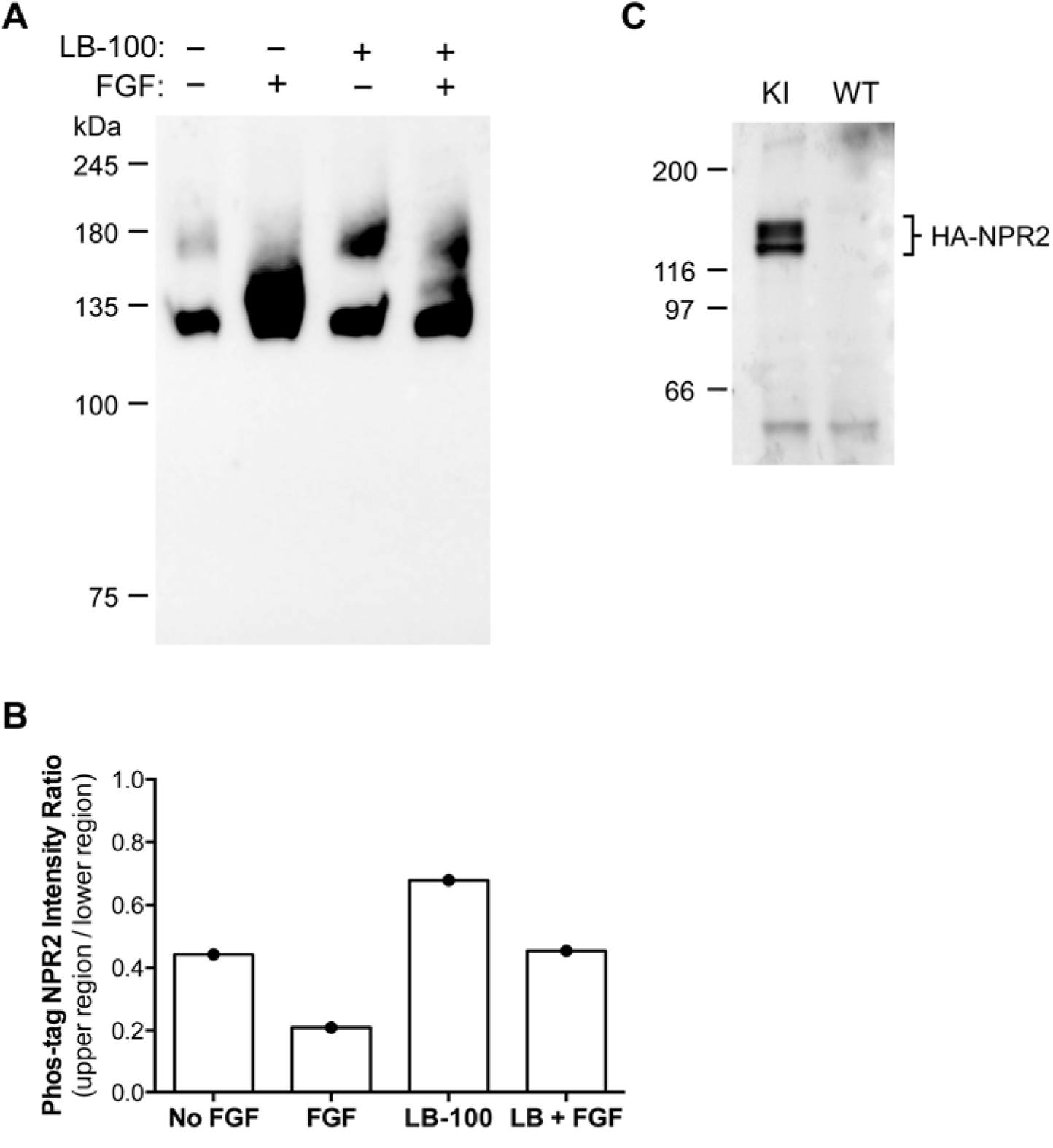
LB-100 counteracts the FGF-induced dephosphorylation of NPR2 in primary chondrocyte cultures. (**A**) Western blot of a Phos-tag gel showing the decrease in NPR2 phosphorylation in chondrocytes treated with FGF18 (0.5 µg/ml) for 10 minutes, and attenuation of this dephosphorylation in chondrocytes pretreated with LB-100 (10 µM) for 1-hour prior to treatment with FGF18. Rib chondrocytes were isolated from mice in which endogenous NPR2 was genetically modified to have an HA epitope tag on its N-terminus (56). The blot was probed with an antibody against HA (Cell Signaling #2367). (**B**) Densitometry measurements from the blot shown in A. The y-axis indicates the ratio of the intensity of the upper region to that of the lower region as shown in ***Figure 2C***; a smaller ratio indicates a decrease in NPR2 phosphorylation. **C**) Western blot of chondrocytes separated on an SDS-PAGE gel without Phos-tag, comparing the chondrocytes from the HA-tagged NPR2 knockin (KI) mouse vs a wildtype (WT) mouse, demonstrating the specificity of the antibody. The 2 bands at ∼130 and ∼120 kDa represent proteins with and without glycosylation (57).

**Figure S3.**
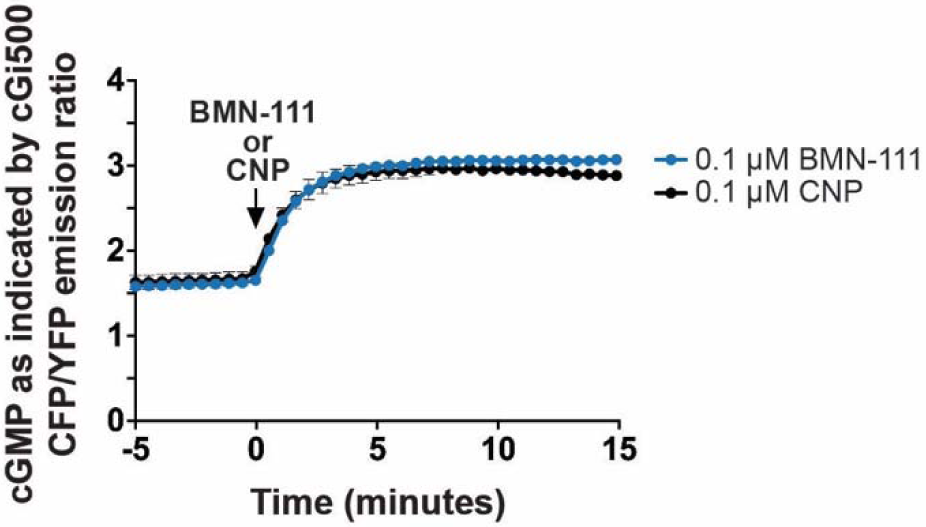
Like CNP, BMN-111 stimulates cGMP production in growth plate chondrocytes. Mean ± SEM for 2 experiments with 0.1 µM BMN-111, and 2 experiments with 0.1 µM CNP, using tibias from littermates.

**Figure S4.**
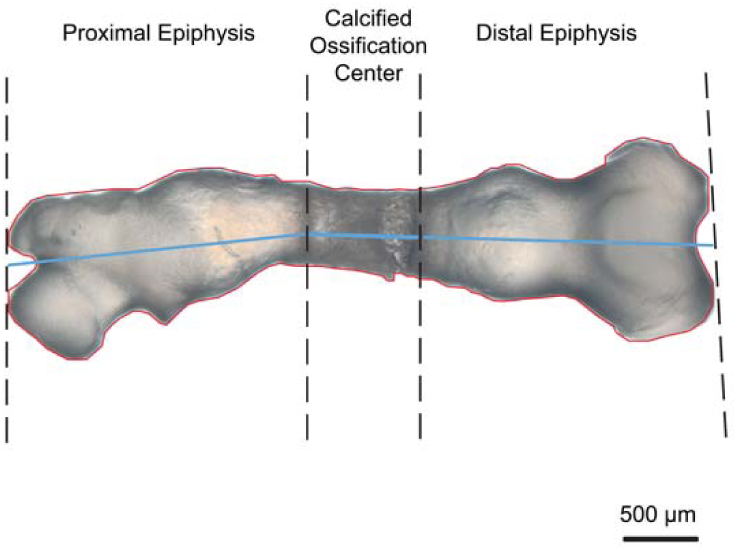
Regions of measurement for bone length and area. The photograph shows a fetal femur from a 16.5 day old *Fgfr3*^*Y367C/+*^ mouse, after 6 days in culture. Bone length was defined as the sum of the lengths of the proximal epiphysis, the calcified ossification center (diaphysis), and the distal epiphysis (blue lines). Bone area was defined as the area within the red line.

**Figure S5.**
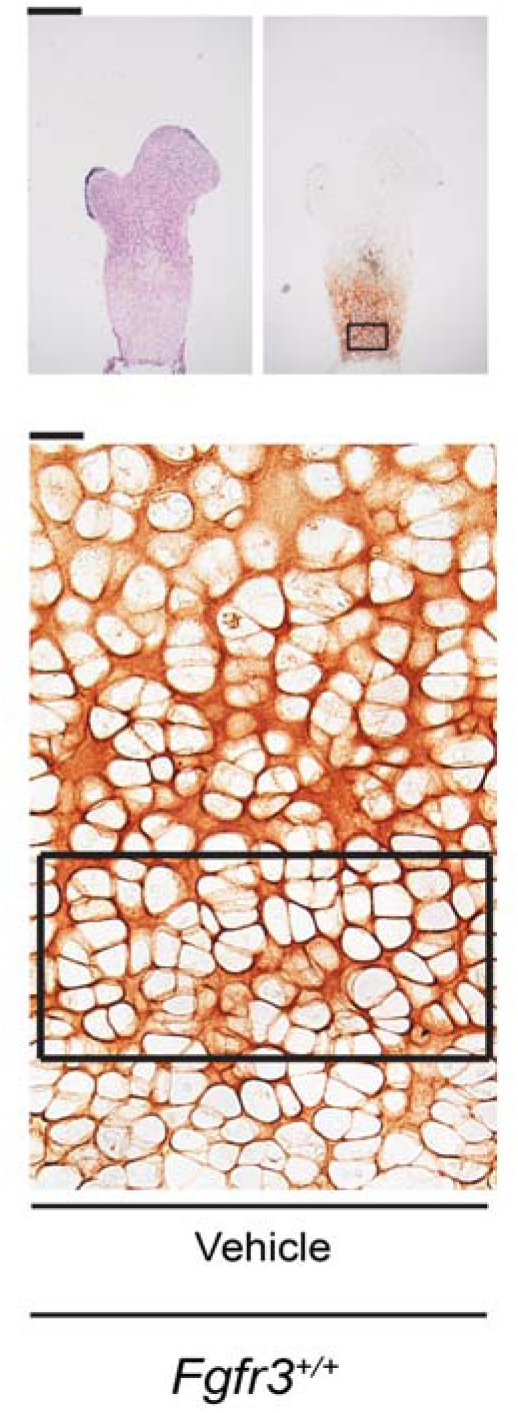
Region used to measure individual chondrocyte areas. A box 166 µm wide ⨯ 76 µm high was positioned at 50 µm above the mineralization front. Images are from ***Figure 4A***.

**Figure S6.**
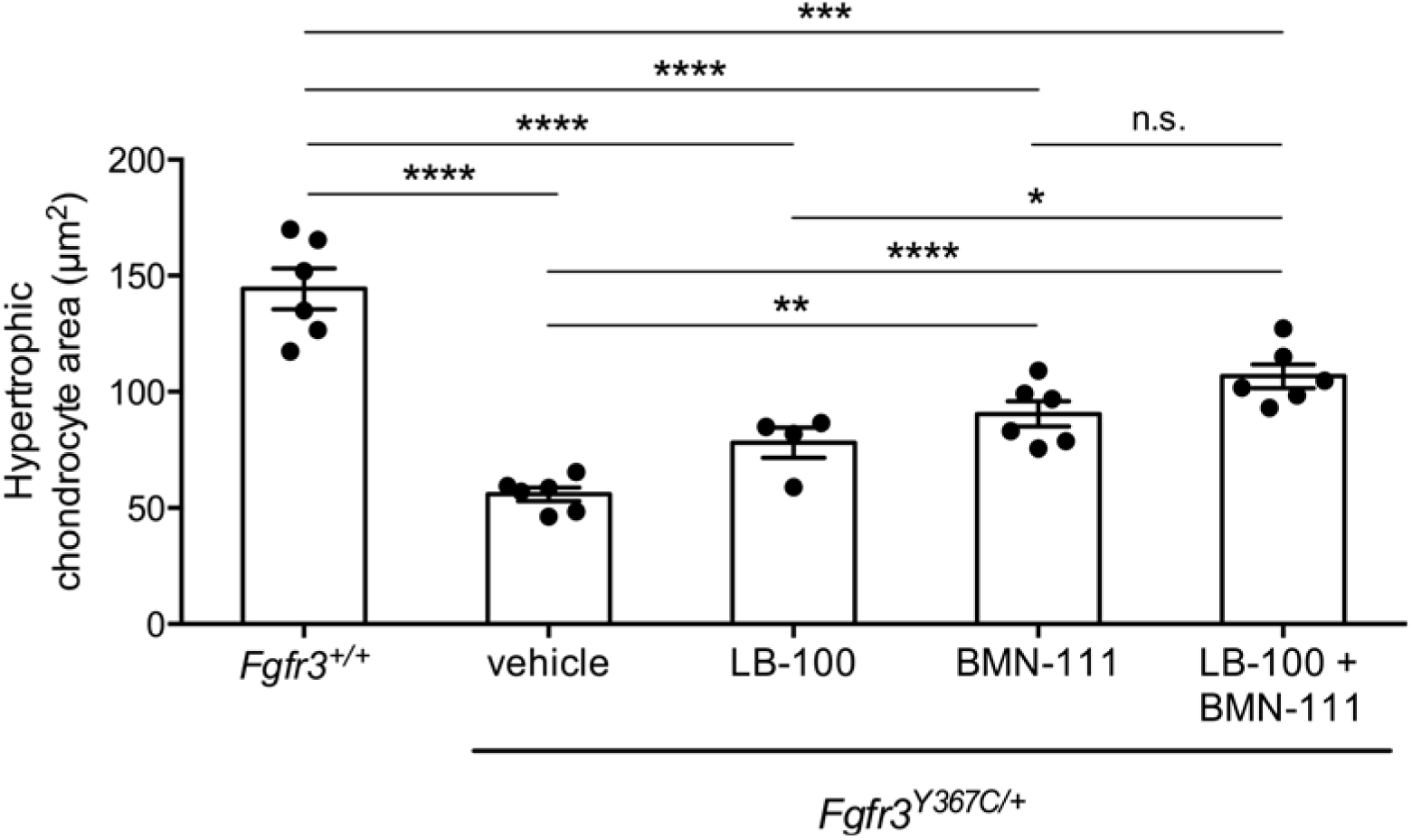
Mean area of individual hypertrophic chondrocytes in distal growth plates of femurs treated as described for the proximal growth plates in ***Figure 4*** (n = 4-6 bones measured for each condition, with 40-134 cells measured for each bone). Data were analyzed by one-way ANOVA followed by the Holm-Sidak multiple comparison test. (*p < 0.05, **p < 0.01, ***p < 0.001, ****p < 0.0001).

**Figure S7.**
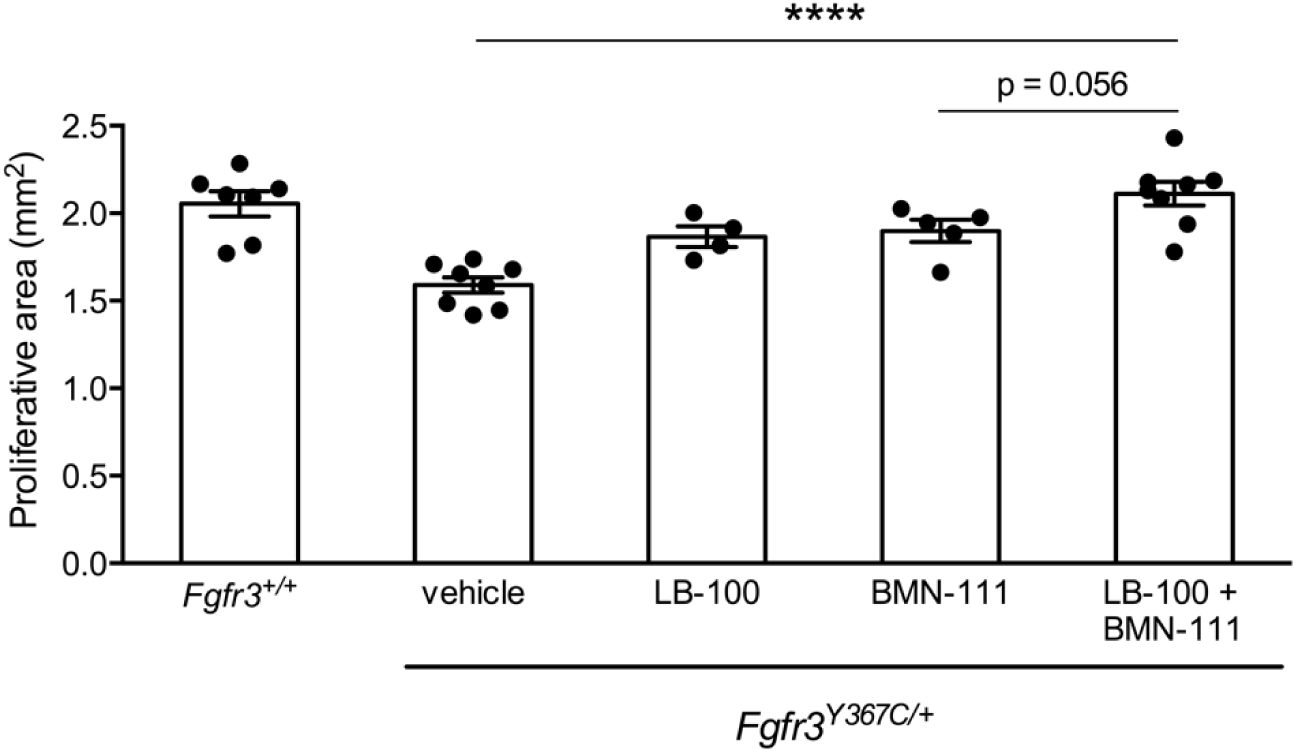
Dual action of LB-100 and BMN-111 improves chondrocyte proliferation in growth plates of ex-vivo cultured *Fgfr3*^*Y367C/+*^ femurs. Total proliferative area of the proximal plus distal growth plates of femurs from *Fgfr3*^*Y367C/+*^ mice, after incubation of the femurs for 6 days with vehicle or LB-100 and/or BMN-111. Growth plates of femurs from *Fgfr3*^*+/+*^) mice cultured with vehicle were also measured. Symbols indicate individual bones (n = 4-8). Bars represent mean ± SEM. Data were analyzed by two-tailed unpaired t-tests between the indicated groups (****p < 0.0001).

